# Impairment Of Aversive Episodic Memories During COVID-19 Pandemic: The Impact Of Emotional Context On Memory Processes

**DOI:** 10.1101/2021.07.06.451294

**Authors:** León Candela Sofía, Bonilla Matías, Urreta Benítez Facundo, Brusco Luis Ignacio, Wang Jingyi, Forcato Cecilia

## Abstract

The threatening context of the COVID-19 pandemic provided a unique setting to study the effects of negative psychological symptoms on memory processes. Episodic memory is an essential function of the human being related to the ability to store and remember experiences and anticipate possible events in the future. Studying this function in this context is crucial to help understand what effects the pandemic will have on the formation of episodic memories. To study this, the formation of episodic memories was evaluated by free recall, recognition, and episode order tasks for an aversive and neutral content. The results indicated that aversive episodic memory is impaired both in the free recall task and in the recognition task. Even the beneficial effect that emotional memory usually has for the episodic order was undermined as there were no differences between the neutral and aversive condition. The present work adds to the evidence that indicates that the level of activation does not modify memory processes in a linear way, which also depends on the type of evocation that people are asked and the characteristics of the content to be encoded.

## 1. Introduction

As COVID-19 cases increased in each region, quarantine was applied in most of the countries as a preventive measure. Thus, because of the different preventive isolation programs the economy was impaired, leading to a strong crisis that critically increased poverty in regions such as South America as well as globally (Buheji et al., 2020) negative impacting in psychological symptoms (Etchevers et al., 2021, Brooks et al., 2020). Specifically, levels of stress, anxiety and depression significantly increased in the population (Salari et al., 2020). In addition, investigations were carried out to determine whether social isolation or ‘‘stay home’’ orders were associated with an increase in negative psychological symptoms (Jacobson et al., 2020). In turn, populations mental health deterioration, translated into an increase in the values of negative mental symptoms such as anxiety defined by Robinson (1990) as ‘‘psychophysiologic sign of stress’’ and depression, can modulate the encoding, consolidation, and retrieval of memories (Cahill &McGaugh, 1998, Beck & Clark, 1997, McGaugh et al., 1996, Roozendaal et al., 2009; beck et al., 2011).

The outbreak of COVID-19 pandemic was considered by several authors as a particularly stressful event due to its novelty, the inability to premeditate and the lack of a vaccine for the cure (Vinkers et al., 2020) as well as it was also thought of as a chronic stressor (Nelson &Bergeman, 2021). In addition, the necessary measures to avoid contagion, (e.g., social isolation), leads to the detriment of people’s mental health, leading to the development of new mental disorders as well as the deepening of already existing ones (Vahia et al., 2020).

Episodic memory is a subtype of declarative memory, defined as the ability to remember where and when past events occurred (Tulving, 1972) and involves subjective consciousness (Wheeler et al., 1997). Furthermore, there is consensus that emotional episodic memories, whether they are pleasant or aversive events, are better remembered than neutral events (Hamann et al., 1999; Payne et al., 2006; Cahill &McGaugh, 1998; Reisberg & Heuer, 2004) proposing that emotional activation mediates episodic memory processes, and it serves as contextual cues for episodic memories (Damasio, 1994, 1999; Bechara et al., 2003; Allen, 2008).

Moreover, emotion improves encoding as well as consolidation processes (LaBar & Cabeza, 2006; Dolcos et al., 2012). Other studies proposed that high emotional arousal could direct attention to prominent details while impairing memory of irrelevant ones (Easterbrook, 1959) and this narrowing of attention may prevent the encoding of relevant information for the memory of events, such as the face of a perpetrator during a crime (Christianson, 1992; Loftus &Messo, 1987).

Both depressive and anxious symptoms can also modulate memory processes (Lachman & Agrigoroaei, 2012; Zlomuzica et al., 2014; Zaheed et al., 2020). In general, it was observed that people with anxiety disorders obtained worse performances than healthy subjects in episodic memory tasks, particularly in the encoding phase, and this impairment is observed even in neutral or irrelevant content (Airaksinen et al., 2005). For the retrieval, the literature generally indicates that this process contains a bias towards negative content (Zlomuzica et al., 2014). On the other hand, the relationship between depression and various cognitive deficits have been widely studied, and it was observed that the impaired domains were working memory, inhibitory control, episodic memory, and speed processing (Airaksinen et al., 2004; Bora, et al., 2013; Gotlib & Joormann, 2010).

The encoding of episodic memories is dependent on the emotional context in which it occurs, and different effects are found according to the content of the material to be learned (Erk et al., 2003). Thus, the COVID-19 pandemic offers a unique opportunity to study in an ecological way how the different types of episodic memory are formed when population are under very demanding psychological conditions,considering that both anxiety and depression are modulating variables of episodic memory function.

While for learning under conditions of previous stress there is broad consensus that this improves emotional content and harms neutral content (Payne et al., 2007), the effect of the before learning stress on consolidation, depending on the type of content, does not seem to be very clear (Shields et al., 2017). Later benefits were found to encoding emotional content under stress (Cahill et al., 2003), although this beneficial effect in the retrieval was also found in both emotional and neutral materials (McCullough & Yonelinas, 2013) and even this effect was observed specifically for neutral information (Yonelinas et al., 2011; Fernández et al., 2015). However, there is a particular work that evaluated the effect of stress generated by a threatening social situation on non-emotional learning and the results indicated that neutral memory was benefited in the short term, and, in addition, the modulation of stress increased the persistence of the memory (Fernández et al., 2015). Similar results in this line were found in paradigms that induced stress by cold pressor Test (CPT) (Schwabe et al., 2008).

We hypothesize that the increase in the values of negative psychological symptoms, such as depressive mood and anxiety, due to the context of the COVID-19 pandemic, modifies the ability to encode and consolidate memories. Thus, we expect that the superiority of emotional episodic memories over neutral ones would be lost not only for the content but also for the temporal order information.

To evaluate this, participants learned an emotional or neutral story on day 1 and were immediately asked for a free recall. Then, on day 8 (one week later) they were asked for a new free recall, a recognition task was carried out and finally a task of the temporal order of events. Then, about 6 months after the first round, the second round was carried out repeating the same procedure but rotating the type of story between the groups.

## 2. Materials and methods

### 2.1. Participants

In total, there were 101 participants, residents of the Metropolitan Area of Buenos Aires, Argentina. They were recruited through advertisements on social networks. Prior to their participation, they signed an informed consent approved by the Alberto Taquini Biomedical Research Ethics Committee. Their ages ranged from 18 to 40 years (25.69 ± 4.36), they did not consume psychotropic drugs and they were not suffering from psychiatric disorders at the time of the experiment. All experiments were carried out virtually. The first round was carried out in April of 2020, within the first month of the period of the Preventive and Mandatory Social Isolation (ASPO) by COVID-19 and the second was carried out in October (6 months later) but within the same quarantine process (Table 1).

**Table 1.**
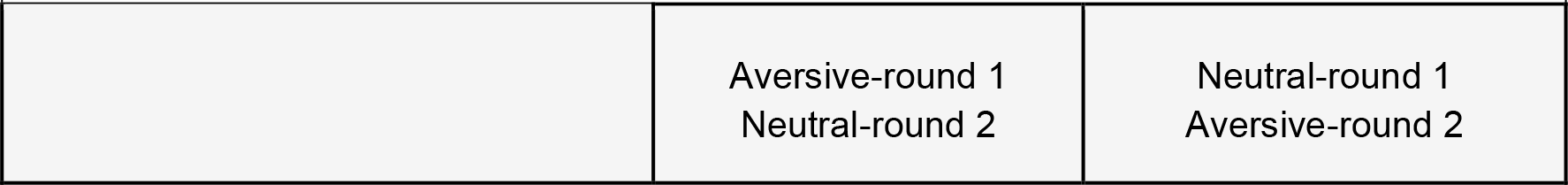
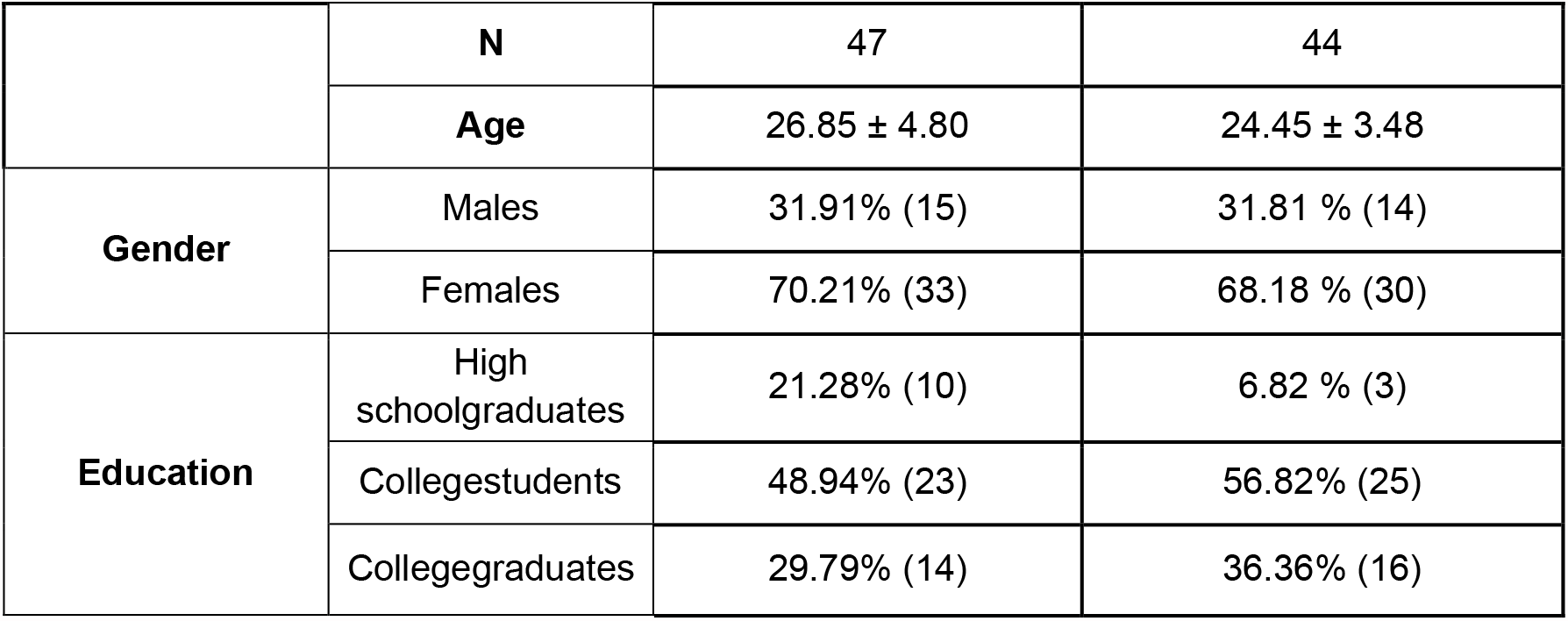
Sociodemographic data. Number of participants in each group, mean age ± SD, percentage of different genders and education level. Participants in the aversive condition of round 1 then became part of the neutral condition in round 2, and vice versa.

| **TABLE 1. Sociodemographic data** |  |  |
| --- | --- | --- |
|  | Aversive-round 1<br>Neutral-round 2 | Neutral-round 1<br>Aversive-round 2 |

|  |  |  |  |
| --- | --- | --- | --- |
|  | **N** | 47 | 44 |
|  | **Age** | 26.85 ± 4.80 | 24.45 ± 3.48 |
| **Gender** | Males | 31.91% (15) | 31.81 % (14) |
|  | Females | 70.21% (33) | 68.18 % (30) |
| **Education** | High schoolgraduates | 21.28% (10) | 6.82 % (3) |
|  | Collegestudents | 48.94% (23) | 56.82% (25) |
|  | Collegegraduates | 29.79% (14) | 36.36% (16) |

### 2.2. Experimental design

The subjects participated in two different rounds of data collection separated by about 6 months. Each participant was randomly assigned to the experimental conditions of the first round. These conditions were formed according to the emotional content of the video that they observed in the training phase, which could be aversive or neutral. The ‘‘Aversive’’ condition was formed of 47 subjects (32 women, 15 men; mean age ± SD: 26.85 ± 4.80) and the neutral condition was formed of 44 people (30 women, 14 men; mean age ± SD: 24.45 ± 3.48). About 6 months later, still within the quarantine period, each participant was assigned to the other experimental condition, e.g., the participants watching the aversive story now watched the neutral one, and the same apply to the other group of subjects. 101 subjects participated in the initial round, 10 of whom decided not to participate in the second round. Therefore, the final number of participants was 91.

### 2.3. Procedure

The experiment was carried out between 10 a.m. and 5 p.m. First, participants received a link via email to complete the sociodemographic questionnaire, the self-administered symptomatology scales: Beck Depression Inventory-II (Beck et al., 1996) and State Trait Anxiety Inventory (Spielberger, 2010). After that, they received a link for the video call and the entire experiment was conducted by one experimenter using this system. Once the connection was established, participants were asked to turn the volume up to maximum and pay attention to the screen. After that, the experimenter’s screen was shared, and the video played. Previously, they were advised that if they had any connection problems during the video, they should notify the experimenter. Nevertheless, this situation did not happen. Immediately after the video, participantsperformed the short-term free recall. One week later, at the same time as day 1, participants entered a video call link previously sent via email. First, they performed the long-term free recall and then the recognition task. It is important to mention that, except when completing the forms, the experimenter was present and available in the video call throughout the experiment. The procedure was repeated in the same way about 6 months later, alternating the content of the stories among the participants (that is, the participants who saw the aversive video later saw the neutral one and vice versa) (Figure 1).

**Figure 1.**
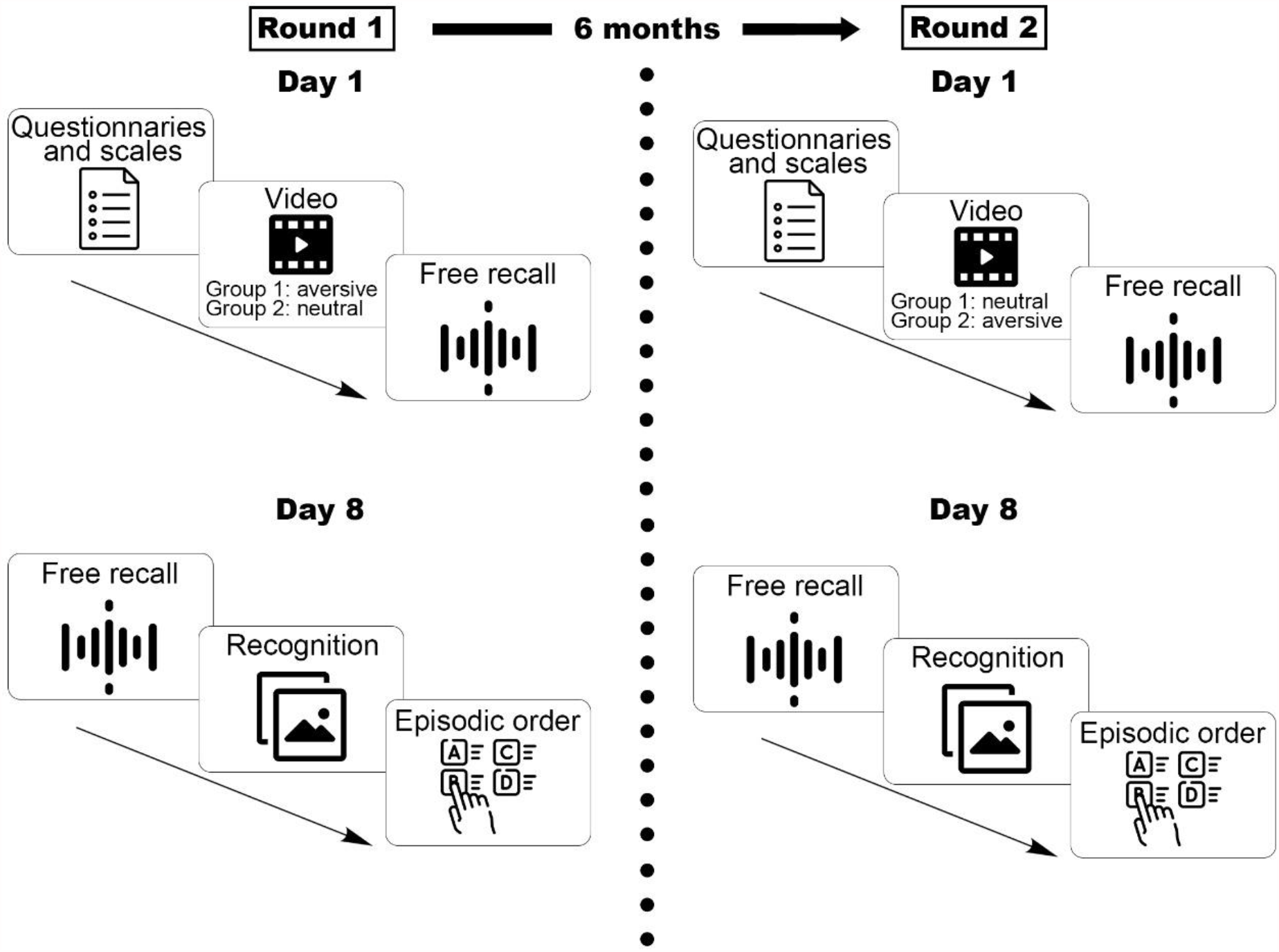
Experimental procedure. The procedure was divided into two parts: First, in round 1 the participants completed day 1 and day 8 both at the same time of day but with a week of difference. One group watched the aversive video, and another group watched the neutral one. After about 6 months, round two was carried out. The procedure was the same, only the content of the videos was alternated. At the end of the experiment, the participants had watched an aversive video and a neutral video with 6 months between one round and the other. Icons ‘‘Choose’’, ‘‘Archivos’’, ‘‘Radio-wave’’, ‘‘Video-player’’ and ‘‘image’’ made by Freepik [https://www.flaticon.com/authors/freepik] from www.flaticon.com

### 2.4. Sociodemographic Questionnaire

This questionnaire consisted of answering questions about age, gender, place of residence during quarantine, chronic medication, and psychiatric disorders.

### 2.5. Symptomatology scales

Self-administered depression scale Beck Depression Inventory-II (Beck et al., 1996) and anxiety scale State Trait Anxiety Inventory (Spielberger, 2010).

### 2.6. Episodic stories

The episodic stories were all consisted of a 3-min video composed of 11 photos each was presented on the screen for 15 seconds, and an auditory narrative describing the on-going story was played during the complete video. The story with aversive emotional content showed a person who suffered from a traffic accident and died in the end. The emotionally neutral story depicted a person’s daily routine. These stimuli were controlled with an independent sample of subjects, who were asked to indicate whether the stimulus was negative, neutral, or positive. 15 additional subjects rated the aversive video and 14 the neutral one. A continuous bar was used that ranged from -100 to -30 for negativity, from -30 to +30 for neutrality and from +30 to +100 for positivity. The participants were asked to choose the value that in their opinion best represented the stimulus (Aversive: -54.00 ± 30.18, Neutral: 11.14 ± 20.91, *t* (27) = 6.70 *p* < 0.001).

### 2.7. Free recall

On Day 1, participants were asked to orally recall the content of the video immediately after watching the video. All oral reportswere recorded, and then the number of correct and false details were counted. Absolute number of reported details about the actions, people, objects, and elements of the environment provided were counted as the subject’s memory performance. Each detail was counted only once, no matter how many times it was repeated in the free recall. The instruction was “Now I am going to ask you to describe to me, in as much detail as possible, what happened in the video. You have no time limit to perform it, do it on your time.’’

The same procedure was conducted again on Day 8 as the delayed recall.

### 2.8. Recognition task

The recognition task consisted of 11 images that were presented altogether, in which, 5 of them were presented in the training session and 6 were new. From these 6 new photos, 3 were similar with the photos presented in this composition, and another 3 were like the images in the video, but not those presented in the composition. Participants had to choose from this series of photos, which of them they considered to have seen in the video. After that, they had to order them according to the episodic order of the story (temporal order). The participants were not limited in time neither to choose the photos they considered correct, nor to order them temporarily. The answer was recorded by an external recorder and then these choices were separated first into correct and incorrect offline. For the temporal order, deviations were accounted for analysis, and this was compared with the correct episodic order of the story. If the participants chose an incorrect photo but placed the event in the correct place in the episodic order, they were taken as valid.

### 2.9. Statistical analysis

These analyzes were carried out in the statistical software IBM SPSS Statistics 25. The scores of the symptomatology scales (STAI and BDI) were used as total values, i.e., the values obtained by the subjects in each test according to the correction rules proposed by each test.

First, with the aim of evaluating whether the values on these scales were elevated in relation to a normal context (before the pandemic), the values obtained by the subjects in both rounds were contrasted with the values of the validations of the symptomatology scales made in the Argentine population by One Sample *t*-test of each of the variables. These validations were carried out in a population like the current sample in terms of place of origin, educational level, age, and gender(Bonicatto et al., 1998; Leibovich de Figueroa, 1991). The values obtained in the symptomatology scales were compared between the two experimental conditions by *t*-test in each round. Also, prior to pooling, it was analyzed that the aversive condition of round 1 does not have differences with the aversive condition of round 2 by *t*-test, and the same was done for the neutral condition. Further, the performance of the conditions in each round was evaluated separately by *t*-test, to control that the results are presented in a similar way at both times point of data collection.

After pooling the subjects, a Paired *t*-test was also carried out to control that there were no differences between the conditions for the values obtained in the symptomatology scales. After confirming this, the pool of subjects was made, and Paired t-tests were performed to compare the performance of the conditions in the memory variables.

As for the free recall details count, it was considered that the number of details that each story presented was not the same, that is, each story provided a different total of possible details to be remembered (Aversive story: 132, Neutral story: 124). Because of this, it was decided to normalize these values using the percentage of true and false details (i.e., N° of correct details/possible correct details*100) and then a memory accuracy index was created by taking the percentage of true details and subtracting the percentage of false details. This calculation was made for both day 1 and 8. In addition, the memory change was calculated for both true details (% true details on day 8 minus % true details on day 1) and for false details (% false details on day 8 less % false details on day 1).

On the other hand, in relation to the recognition task, both the number of correct and incorrect choices were calculated. Regarding the episodic order task, the number of deviations with respect to the correct temporal order was registered. For all these analyses, a paired *t-*test was performed to compare the performance of the conditions.

Finally, to examine whether the encoding and consolidation processes could be related by negative psychological symptoms, we analyzed the values obtained on the symptomatology scales and the performance obtained in each task by both conditions with Pearson correlations. Alpha was set at 0.05, we did not perform corrections for multiple comparisons.

## 3. Results

### 3.1. Symptomatology scales

Regarding the levels of state anxiety, trait anxiety and depression of the aversive and neutral conditions we found a significant increase during the quarantine by COVID-19 compared to the mean level of the Argentinean population (mean values of the validation scales) before the quarantine (Table 1. Supplementary material). We further found no significant differences for the symptomatology scales neither between the two rounds of data collection (month 1 and month 6) (Table 2. Supplementary material) nor between the rounds in each condition (neutral or aversive content) for these same variables (Table 3. Supplementary material). Finally, as there were no significant differences neither in any of the memory variables studied here between rounds (Table 4. Supplementary material) nor between the rounds of each condition (Table 5. Supplementary material) we pooled the data of each condition. Taken together, these results indicate that the values obtained in the symptomatology scales were higher at the first month of quarantine than the norm, and it remained high after about 6 months. In the same way, it is observed that the different types of information are affected in the same way at both periods of data collection.

### 3.2. Symptomatology scales after pooling

We pooled the data of the two rounds according to the emotional content of the learned story, observing no significant differences between the conditions for the values obtained in the symptomatology scales (Table 2).

**Table 2.**
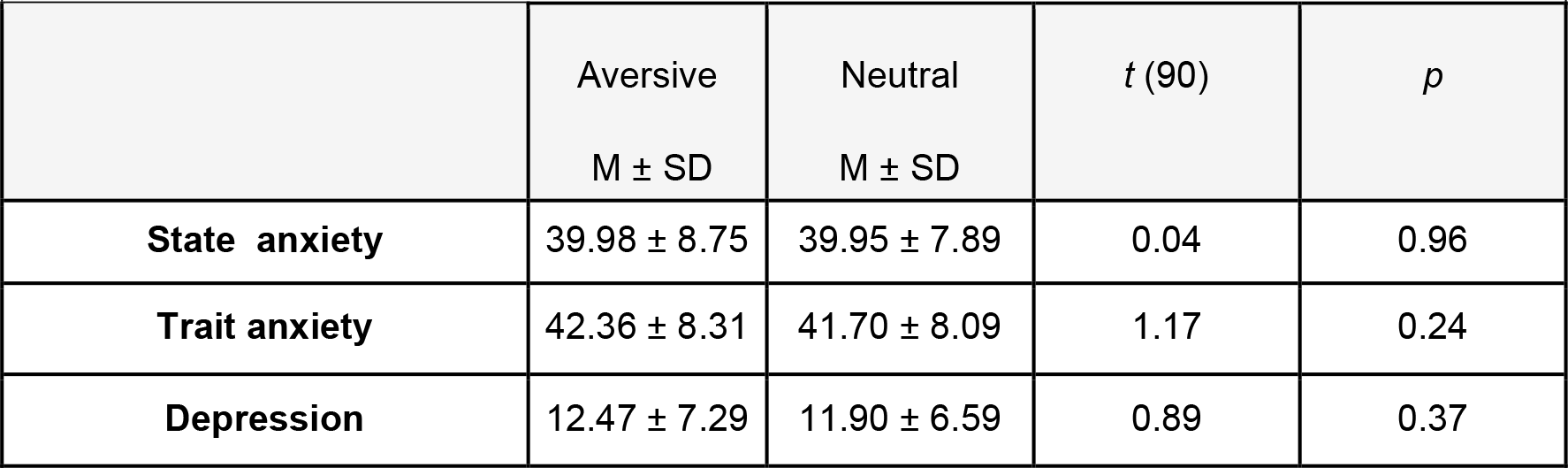
Symptomatology scales after pooling. Mean state anxiety, trait anxiety, depression ± SD. A Paired t-test was used to compare symptom scale scores between the conditions Aversive and neutral.

### 3.3. Episodic memory tasks

First, we found that the neutral condition reached a significantly higher accuracy index (percentage of true details minus the percentage of false details) in the free recall task on day 1 (Aversive: 33.84 ± 9.43, Neutral: 40.09 ± 11.07, *t* (90) = -5.32 *p* < 0.01) and on day 8 (Aversive: 24.04 ± 9.06, Neutral: 30.10 ± 11.06, *t* (90) = -5.60 *p* < 0.01). However, when we analyzed the memory change for true details (% true details day 8 minus % true details day 1), no significant differences between conditions were found (Aversive: -24.67 ± 16.71, Neutral: -19.59 ± 26.96, t (90) = - 1.75 *p* = 0.08) neither for false details (Aversive: 1.07 ± 1.74, Neutral: 1.07 ± 2.03, *t* (90) = 0.01 *p* = 0.99) (Figure 2).

**Figure 2.**
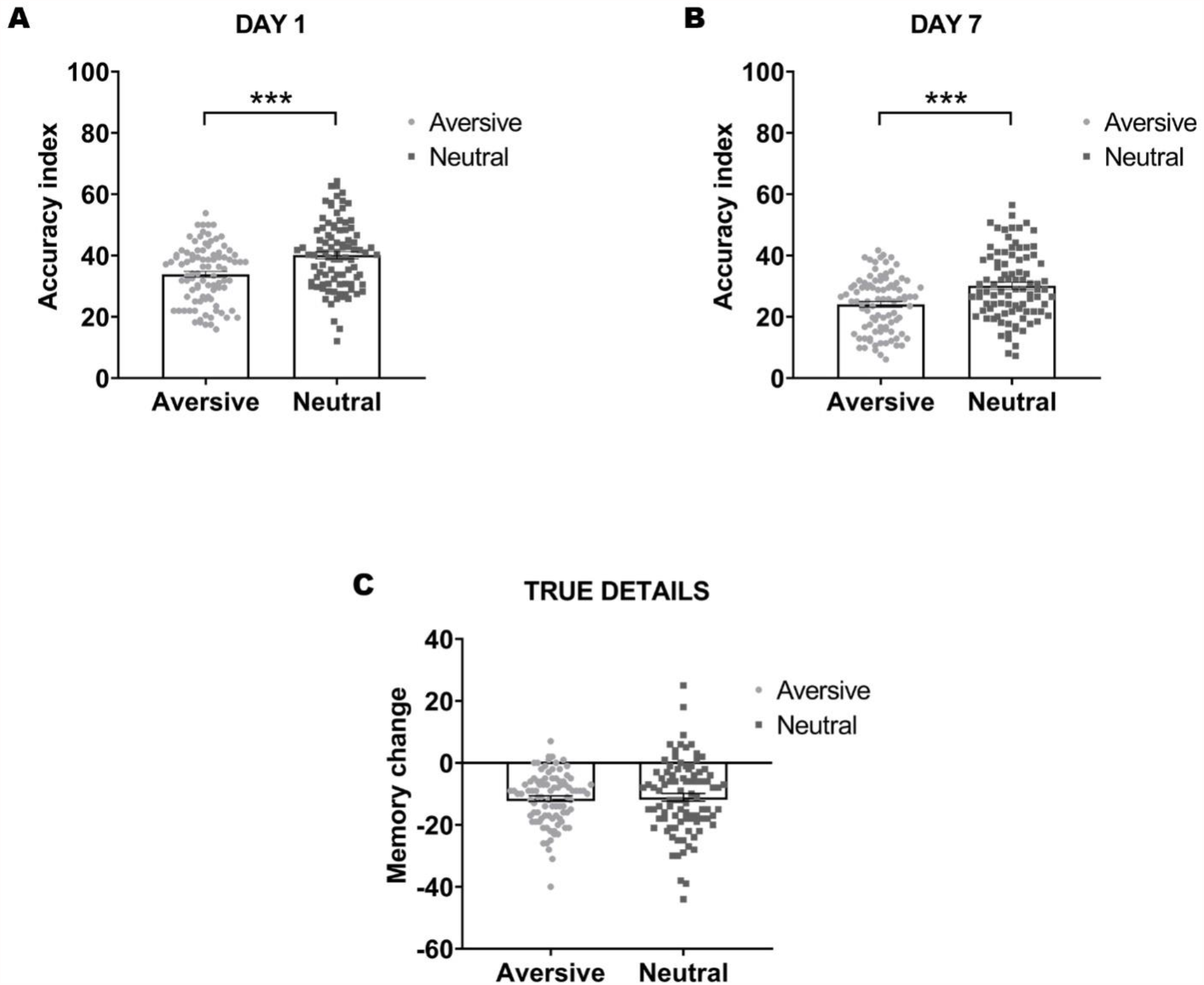
Free recall. This figure indicates the performance of the conditions in the free recall task. **A**. M ± SEM Accuracy index of day 1 (% true details less % false details). **B**. M ± SEM Accuracy index of day 8 (% true details less% false details). **C**. Memory change between day 1 and day 8 regarding the true details (% of true details at day 8 less % of true details day 1)

Regarding the recognition task, the neutral condition showed significantly higher number of correct choices than the aversive condition (Aversive: 3.93 ± 1.08, Neutral: 4.24 ± 0.83, *t* (90) = -2.37 *p* = 0.02), but they did not differ in the number of incorrect the choices (Aversive: 0.84 ± 1.04, Neutral: 0.77 ± 0.92, *t* (90) = 0.61 *p* = 0.54). Moreover, no differences were observed between the conditions in terms of the number of deviations from the correct episodic order (Aversive: 1.23 ± 1.27, Neutral: 1.18 ± 1.22, *t* (90) = 0.32 *p* = 0.74) (Figure 3).

**Figure 3.**
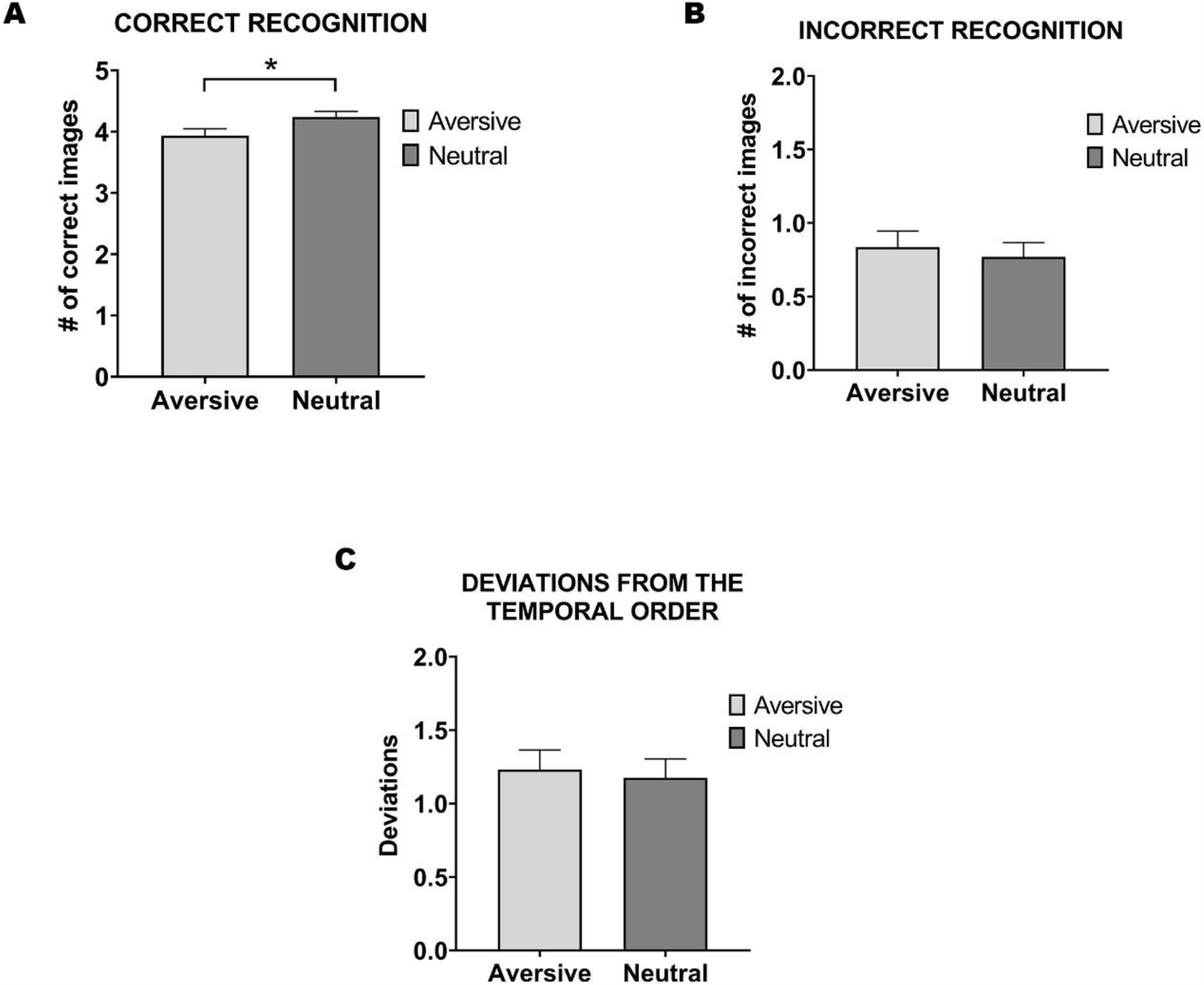
Recognition and episodic order. This figure indicates the performance of the conditions in the recognition and episodic order task. **A**. M ± SD correct images chosen. **B**. M ± SD incorrect images chosen. **C**. M ± SD deviations from the correct episodic order.

### 3.4. Symptomatology Scales and free recall

Since negative psychological symptoms modulate the memory processes, we further performed correlations analyses between the symptomatology scales values obtained and the values obtained in the different free recalls (day 1 and day 8) of each condition. We first found a significant negative correlation between state anxiety and the accuracy index on day 1 for the aversive condition (r = -0.32, *p* = 0.002), while this result was not observed in the neutral one (r = -0.12, *p* = 0.25). We further observed significant negative correlations between these same variables for both conditions on day 8 (Aversive: r = -0.22, *p* = 0.02; Neutral: r = -0.25, *p* = 0.01). Regarding depression, we found negative correlations between this variable and the accuracy index on day 8 for both conditions, but it was only significant for the neutral one (Aversive: r = -0.16 *p* = 0.13; Neutral: r = -0.21, *p* = 0.04) (Figure 4). These results indicate that the higher the anxiety on day 1, the worse the performance in learning aversive content. In this way, it seems that the encoding of aversive content is more affected by the state anxiety than the neutral one, while consolidation and evocation seem to be affected in the same way.

**Figure 4.**
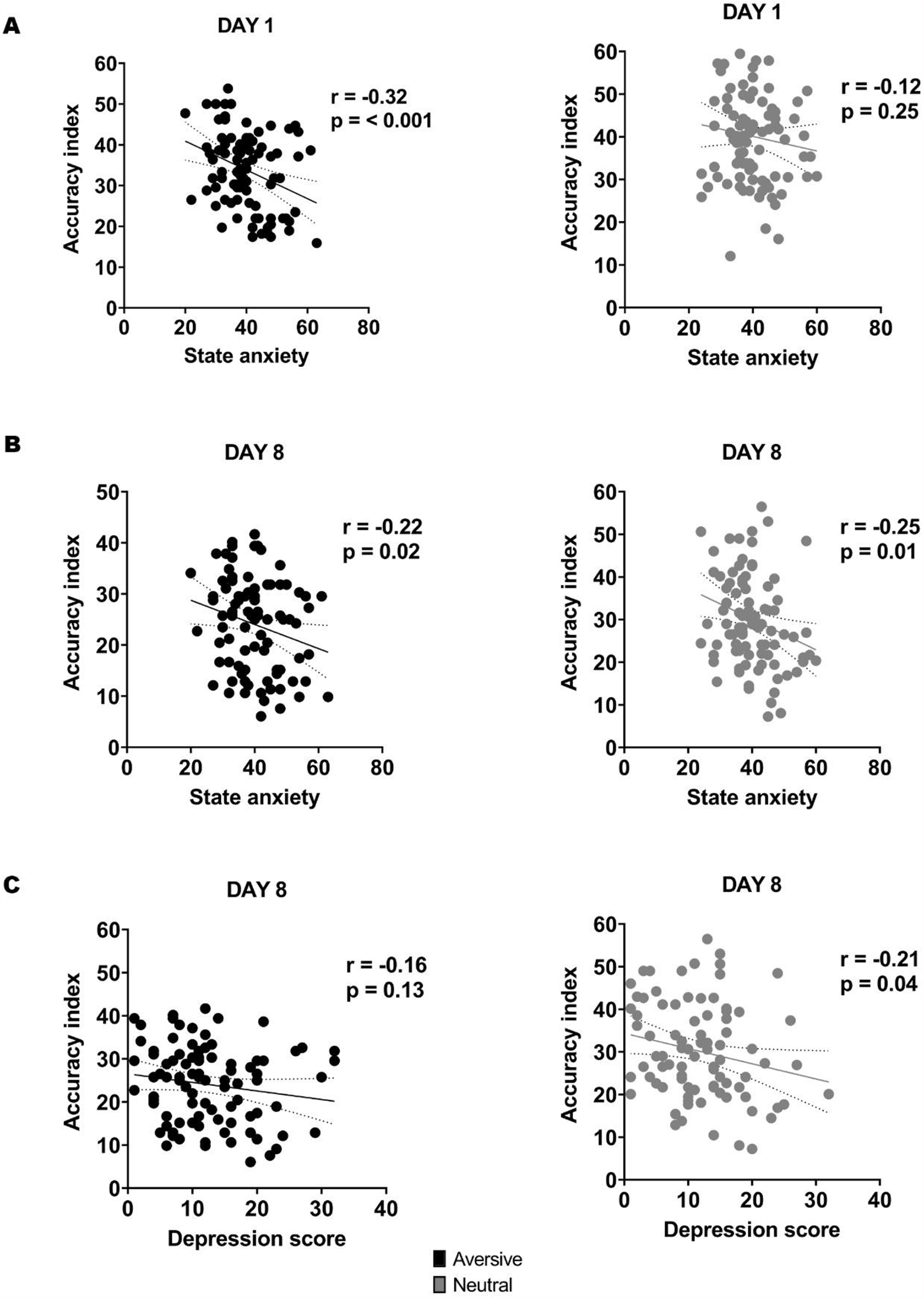
Symptomatologic scores and episodic memory performance. **A**. Correlations of both conditions between state anxiety and accuracy index day 1. **B**. Correlations of both conditions between state anxiety and accuracy index day 8. **C**. Correlations of both conditions between depression and accuracy index day 8.

### 3.5. Symptomatology Scales and recognition task

We then evaluated the relationship between the values obtained in the symptomatology scales and the measures obtained both in the recognition task and episodic order task. We found a significant positive correlation between the values obtained on the depression scale and the number of incorrect choices for the aversive condition (r = 0.22, *p* = 0.03). However, no significant correlation was found for the neutral condition (r = -0.11, *p* = 0.27). No other significant correlations were found (-0.16 < r < 0.20, all *ps* > 0.05).

## 4. Discussion

Primarily, we observed that the levels of anxiety and depression were significantly increased with respect to the population mean of a normal context. These results coincide with large-scale surveys of the Argentine population during the pandemic, which indicate a significant increase in anxiety and depressive symptoms even in the early stages of the pandemic, in people who had never presented these symptoms (Torrente et al., 2020). It is interesting to propose that the pandemic *per se* might not serve as a unified source of stress, rather, different sources of stressors may co-exist within this phenomenon of the COVID-19 pandemic (Biondi & Iannitelli, 2020): On the one hand, the existence of a virus that may eventually be the cause of death of our own or our loved ones, on the other hand ‘‘ infodemic ‘‘ a term coined by the World Health Organization to define the distressing excess of information (Zarocostas 2020), and finally the isolation and lockdown measures that led to a total disturbance of people’s social routine (Galea et al, 2020). In sum, we propose that Argentina, particularly the metropolitan area where the epidemiological situation was the most serious, offered a unique opportunity to study the influence of these stressors together.

The main objective of this work was to analyze the effects of the threatening COVID-19 pandemic on the formation and consolidation of episodic memory. We were particularly interested in understanding how main changes of people’s mental health in the pandemic (e.g., anxiety and depression that are known as modulators of episodic memory (Airaksinen et al., 2004; 2005; Gotlib & Joormann, 2010; Zlomuzica et al., 2014) might modulate our memories.

Due to the threatening context, we expected that the superiority of emotional episodic memories over neutral ones would be lost for the content. However, contrary to our expectations, we found that this situation particularly impacted on the processing of memories with emotional content, since in addition to losing the memory enhancing effect, the aversive content was remembered significantly less than the neutral.

Our results indicated that the neutral condition obtained a better score in the accuracy index in both the short-term free recall and the long-term free recall. We suggested that the difference between conditions is related to the encoding process and not to consolidation because no differences between conditions were found for the memory change. In addition, neutral content was recognized better than aversive and contrary to previous results that indicate that temporal order is favored by emotion compared to non-emotional content (Groch et al., 2011), we found that this difference is missing. Thus, we suggest that the pandemic context, translated into increases in anxiety, depression could be impairing the encoding of aversive content.

However, the general literature indicates that high levels of stress improve emotional memory encoding (Buchanan and Lovallo, 2001; Cahill et al., 2003; McGaugh, 2000). Although, it is necessary to highlight that in this group of studies the stress was induced with cortisol administration. Instead, in the population chosen for this work, immersed in one of the most shocking global events in recent years, the activation is generated by the context of life (Galea et al., 2020). On the other hand, there are works in which people were exposed to psychological stress through an exposure to the public task prior to learning and it was found that the emotional state impaired the memory of neutral words reducing the number of details provided (Kirschbaum et al., 1996). One of the possible explanations for these apparently contradictory results could be the Yerkes-Dodson curve (Yerkes & Dodson, 1908). This phenomenon has broad evidence, and it states that there is an optimal point of stress where the cognitive performance seems to be strengthened, whereas if the stress is too low or too high this effect disappears. Moreover, if the stress level is too high (e.g., by strong stress or anxiety) the memory recall performance is impaired. In our results, the COVID-19 pandemic could have caused high levels of stress impairing the aversive content, leading to not only losing the beneficial effect of emotion but also being harmed compared to non-emotional content (Yerkes & Dodson, 1908; Mair et al., 2010).

Two main brain regions have been proposed to be involved in the emotion enhancement effect in episodic memory (Maratos et al., 2001): areas of the amygdala and hippocampus that contribute to the encoding, consolidation, and retrieval of emotional memories (Kensinger & Corkin, 2004; McGaugh, 2004; Ritchey et al., 2008; Kensinger & Schacter, 2005) and prefrontal cortex that benefits executive, attentional, and semantic processes (Dolcos et al., 2017). However, it has been proposed that at the high end of the Yerkes Dodson curve, where the individual is suffering high level of stress, the function of PFC could be suppressed for excessive emotionality (Diamond et al., 2007). In addition, the involvement or not of the prefrontal cortex has been thought as a delimiter between ‘‘ simple ‘‘ and ‘‘ complex ‘‘ tasks, and in turn it has been observed that memory tasks that present the curvilinear relationship of the Yerkes Dodson curve between activation and performance are precisely those that involve this brain area (Diamond et al., 2007). Also, in lesion and neuroimaging studies of the prefrontal cortex, complex tasks such as those involving free recall without cues were particularly affected compared withsimple recognition tasks that involve yes-no responses (Ranganath et al., 2003; Ranganath & Knight 2005). Taken together, we propose that the increased levels of anxiety and depression due to the COVID-19 pandemic could have increased the stress from its optimal level, hence impairing the encoding of new episodic memory.

As previously noted, stress distinctively affects types of retrieval: while free recall tasks can be impaired, recognition tasks may be not modified, and this is due to differences in difficulty related to the availability or not of retrieval cues (Gagnon & Wagner, 2016). However, our results indicate that although the aversive condition did not present more incorrect choices, it chose significantly fewer correct images than the neutral condition. This could be indicating a comprehensive modification in the way of encoding aversive memories that affects both types of recall, regardless of the difficulty of the task. Nevertheless, more studies are needed to compare both types of recall in the context of a pandemic depending on the type of memory that is learned to expand the knowledge in this area. Even in the future, the effect of remembering traumatic memories specifically related to the situation of the COVID-19 pandemiccan be studied.

Additionally, the fact that neutral content is better encoded than aversivecould be explained by the possible lack of novelty of the aversive stimulus, that is, receiving information about an accident that produces a death may be encoded as a non-novel memory when the news about deaths or possible deaths are present constantly in everyday life. In this way, it could be relatively easy to process this type of information due to its familiarity component (Waris et al., 2021).

The main limitation of this work is the fact of not being able to compare the measurements obtained with measurements outside the context of the pandemic. In Argentina we are still going through the “second wave” of the pandemic and living conditions are far from normal. Finally, it would have been desirable to have a physiological measure of people’s stress (for example blood cortisol measurements), but this was impossible due to the obligation to carry out the experiments virtually. However, we obtained measures of anxiety, previously defined as the psychophysiological signal that the stress response has been initiated (Robinson, 1990). Last but not the least, we did not have a post-event emotional rating scalewhich indicates the emotional state caused by the stimulus. However, the negativity or neutrality of the stories was controlled with an independent group of participants and both stories served their purpose.

First, our results add to the increasing evidence that human basic cognitive function is negatively affected by the COVID-19 pandemic context. Second, we provided novel evidence that episodic memory attached with different emotional content are affected differently from normal context. Last but not the least, taking into account that episodic memory is the ability to recall information about what happened, where it happened, and when the event happened (Tulving, 1993) and is a central function for the everyday life, our study can provide information for future psychological treatments that address, for example, Post-Traumatic Stress Disorders (PTSD) related to the pandemic that are already foreseen (Kathirvel, 2020; Carmassi et al., 2020), or simply to understand how the formation of memory of events works in highly demanding environments.

## 6. Author Contributions

CSL, MB, FAUB, JW and CF made substantial contributions to the conception and design of the work. CSL and MB ran the experiments. CSL and MB performed the statistical analyses. CSL and CF contributed by drafting the work. CLS, MB, FAUB, BIL, JW and CF contributed to revising it critically.

## Supplementary Material

**Table 1.**
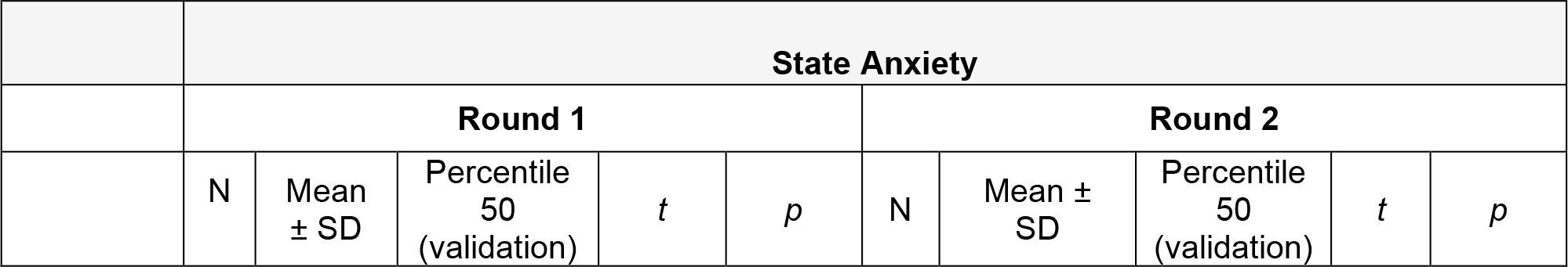
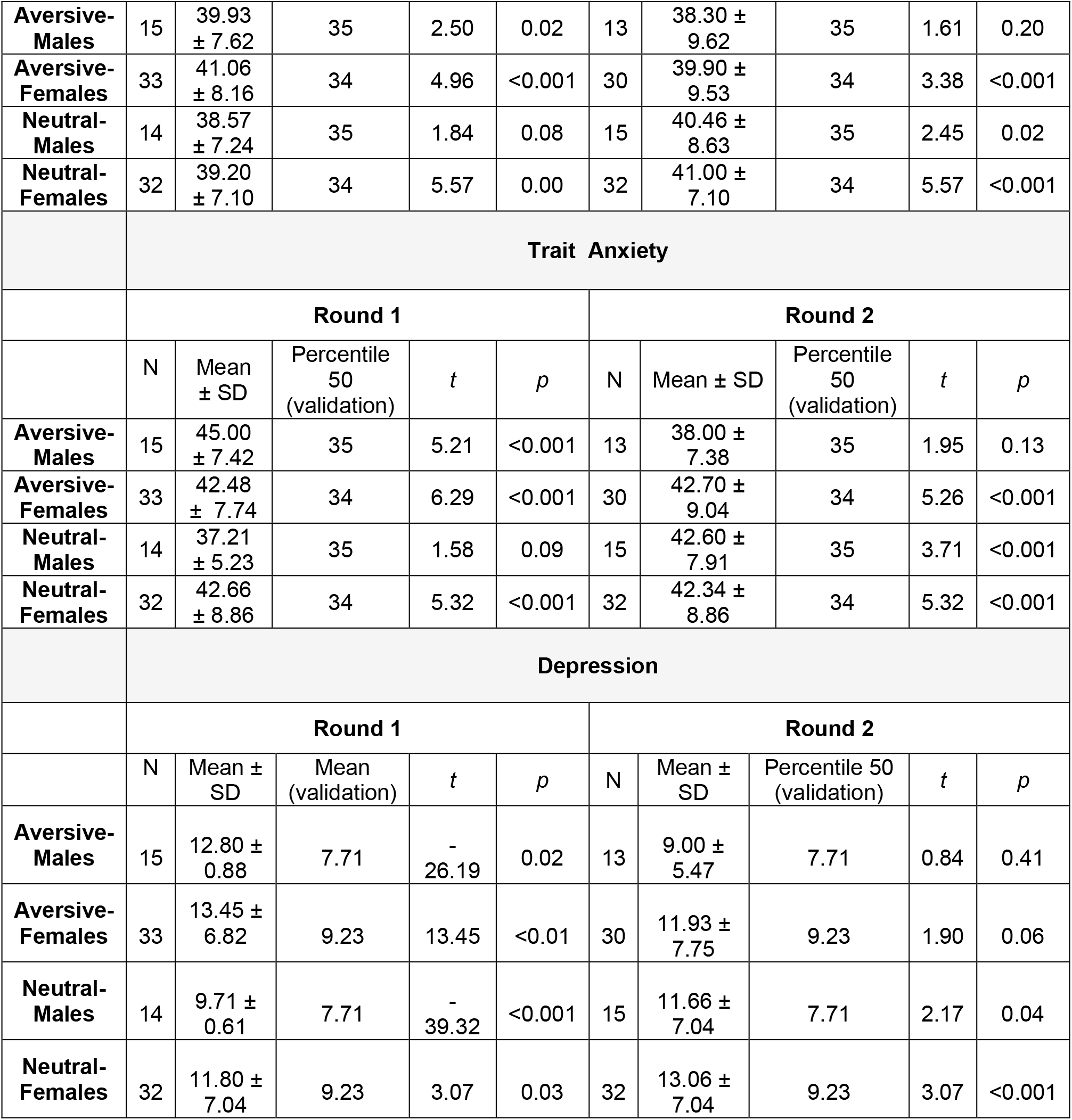
Symptomatology scales. Comparison between the present sample and the mean level of the Argentinean population before the quarantine for state anxiety, trait anxiety and depression. A two-tailed t-test was used to compare symptom scale scores between the conditions Aversive and Neutral and the validation values. The separation between males and females was made because this is done in the validations.

**Table 2.** Symptomatology scales. Comparison between the conditions of each round. Mean state anxiety, trait anxiety, depression ± SD. A two-tailed t-test was used to compare symptom scale scores between the Aversive/Neutral conditions and the validation values. The separation between males and females was made because this is done in the validations.

| TABLE 2. Symptomatology scales |  |  |  |  |  |  |  |  |
| --- | --- | --- | --- | --- | --- | --- | --- | --- |
|  | Round 1 |  |  |  | Round 2 |  |  |  |
| | Aversive<br>M ± SD | Neutral<br>M ± SD | *t* (89) | *p* | Aversive<br>M ± SD | Neutral<br>M ± SD | *t* (89) | *p* |
| **State anxiety** | 40.70 ±<br>8.12 | 38.18 ±<br>10.03 | 0.95 | 0.34 | 39.34 ±<br>9.43 | 40.82 ±<br>7.53 | 0.87 | 0.38 |
| **Trait anxiety** | 43.31 ±<br>8.00 | 39.93 ±<br>9.61 | 1.63 | 0.10 | 40.27 ±<br>10.08 | 42.42 ±<br>8.48 | 0.75 | 0.45 |
| **Depression** | 13.76 ±<br>7.28 | 10.95 ±<br>6.29 | 1.89 | 0.07 | 10.95 ±<br>7.18 | 12.61 ±<br>7.00 | 1.06 | 0.29 |

**Table 3.** Symptomatology scales. Comparison between rounds of each condition. Mean state anxiety, trait anxiety, depression ± SD. A two-tailed t-test was used to compare symptom scale scores between the rounds of each condition before pooling subjects.

| TABLE 3. Symptomatology scales |  |  |  |  |  |  |  |  |
| --- | --- | --- | --- | --- | --- | --- | --- | --- |
|  | Aversive |  |  |  | Neutral |  |  |  |
| | Round 1<br>M ± SD | Round 2<br>M ± SD | *t* (89) | *p* | Round 1<br>M ± SD | Round 2<br>M ± SD | *t* (89) | *p* |
| **State anxiety** | 40.70 ±<br>8.12 | 39.34 ±<br>9.43 | 0.74 | 0.46 | 38.18 ±<br>10.03 | 40.82 ±<br>7.53 | -1.10 | 0.27 |
| **Trait anxiety** | 43.31 ±<br>8.00 | 40.27 ±<br>10.08 | 1.44 | 0.15 | 39.93 ±<br>9.61 | 42.42 ±<br>8.48 | -0.87 | 0.38 |
| **Depression** | 13.76 ±<br>7.28 | 10.95 ±<br>7.18 | 1.82 | 0.07 | 10.95 ±<br>6.29 | 12.61 ±<br>7.00 | -1.07 | 0.28 |

**Table 4.** Memory tasks. Comparison between conditions of each round. Mean memory variables ± SD. A two-tailed t-test was used to compare the performance at memory tasks between the conditions of each round before pooling subjects.

| TABLE 4. Memory tasks |  |  |
| --- | --- | --- |
|  | Round 1 | Round 2 |

| | Aversive<br>M ± SD | Neutral<br>M ± SD | *t* (89) | *p* | Aversive<br>M ± SD | Neutral<br>M ± SD | *t* (89) | *p* |
| --- | --- | --- | --- | --- | --- | --- | --- | --- |
| **Accuracy Index (day 1)** | 32.75 ± 10.09 | 39.30 ± 10.93 | -2.97 | 0.004 | 35.02 ± 8.63 | 40.84 ± 11.26 | 2.75 | 0.007 |
| **Accuracy Index (day 8)** | 22.84 ± 9.33 | 30.75 ± 10.04 | -3.89 | <0.001 | 25.32 ± 8.68 | 29.49 ± 12.02 | 1.88 | 0.06 |
| **Memory change (True details %)** | -25.88 ± 17.24 | -13.41 ± 27.60 | -2.51 | 0.01 | -25.10 ± 25.54 | -23.54 ± 16.18 | -0.34 | 0.73 |
| **Memory change (False details %)** | 1.27 ± 1.68 | 1.66 ± 2.80 | -0.87 | 0.38 | 0.46 ± 1.28 | 0.79 ± 1.78 | -1.02 | 0.31 |
| **Correct choices** | 3.91 ± 1.03 | 4.09 ± 0.91 | -0.85 | 0.39 | 3.95 ± 1.14 | 4.38 ± 0.73 | 2.14 | 0.03 |
| **Incorrect choices** | 0.98 ± 1.09 | 0.59 ± 0.78 | 1.93 | 0.05 | 0.68 ± 0.98 | 0.94 ± 1.00 | 1.21 | 0.22 |
| **Deviations** | 1.15 ± 1.16 | 1.30 ± 1.23 | -0.58 | 0.56 | 1.32 ± 1.39 | 1.06 ± 1.22 | -0.92 | 0.35 |

**Table 5.**
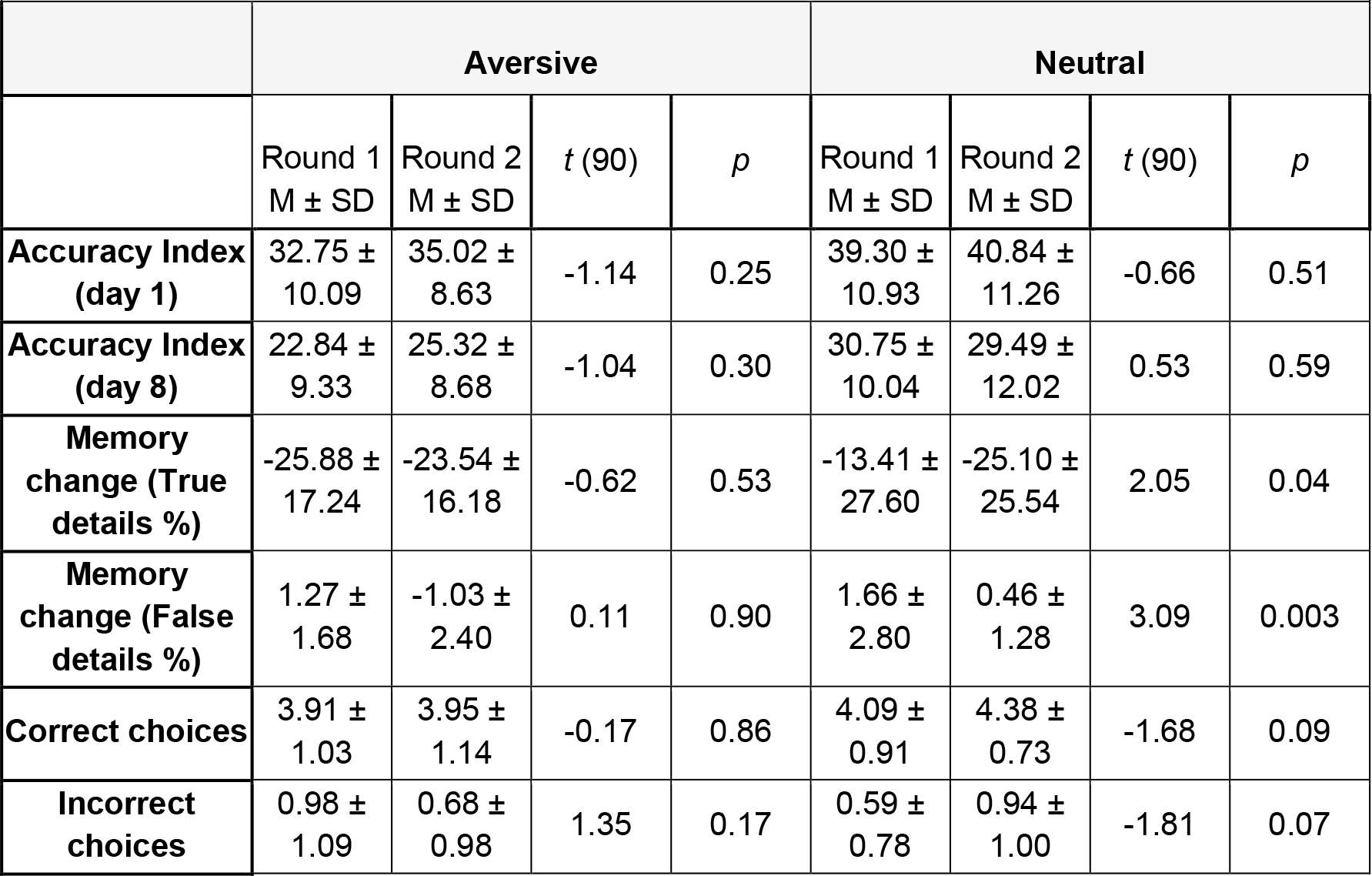
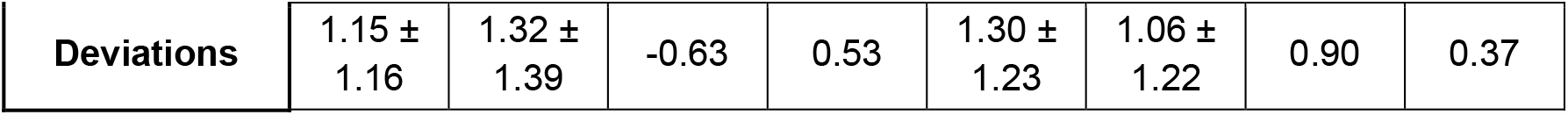
Comparison between rounds of each condition. Mean memory variables ± SD. A two-tailed t-test was used to compare the performance at memory tasks between the rounds of each condition before pooling subjects.

| **TABLE 5. Memory tasks** |  |  |  |  |  |  |  |  |
| --- | --- | --- | --- | --- | --- | --- | --- | --- |
|  | **Aversive** |  |  |  | **Neutral** |  |  |  |
| | Round 1<br>M ± SD | Round 2<br>M ± SD | *t* (90) | *p* | Round 1<br>M ± SD | Round 2<br>M ± SD | *t* (90) | *p* |
| **Accuracy Index (day 1)** | 32.75 ± 10.09 | 35.02 ± 8.63 | -1.14 | 0.25 | 39.30 ± 10.93 | 40.84 ± 11.26 | -0.66 | 0.51 |
| **Accuracy Index (day 8)** | 22.84 ± 9.33 | 25.32 ± 8.68 | -1.04 | 0.30 | 30.75 ± 10.04 | 29.49 ± 12.02 | 0.53 | 0.59 |
| **Memory change (True details %)** | -25.88 ± 17.24 | -23.54 ± 16.18 | -0.62 | 0.53 | -13.41 ± 27.60 | -25.10 ± 25.54 | 2.05 | 0.04 |
| **Memory change (False details %)** | 1.27 ± 1.68 | -1.03 ± 2.40 | 0.11 | 0.90 | 1.66 ± 2.80 | 0.46 ± 1.28 | 3.09 | 0.003 |
| **Correct choices** | 3.91 ± 1.03 | 3.95 ± 1.14 | -0.17 | 0.86 | 4.09 ± 0.91 | 4.38 ± 0.73 | -1.68 | 0.09 |
| **Incorrect choices** | 0.98 ± 1.09 | 0.68 ± 0.98 | 1.35 | 0.17 | 0.59 ± 0.78 | 0.94 ± 1.00 | -1.81 | 0.07 |
| **Deviations** | 1.15 ±<br>1.16 | 1.32 ±<br>1.39 | -0.63 | 0.53 | 1.30 ±<br>1.23 | 1.06 ±<br>1.22 | 0.90 | 0.37 |

